# Differential biological effect of low doses of ionizing radiation depending on the radiosensitivity in a cell line model

**DOI:** 10.1101/2024.05.22.595283

**Authors:** Elia Palma-Rojo, Joan-Francesc Barquinero, Jaime Pérez-Alija, Juan R González, Gemma Armengol

## Abstract

**Purpose:** Exposure to low doses (LD) of ionizing radiation (IR), such as the ones employed in computed tomography (CT) examination, can be associated with cancer risk. However, not all individuals respond the same to IR, and cancer development could depend on the individual radiosensitivity. Notably, inter-individual differences in the response to IR have been very well studied for high and medium doses, but not for LD. In the present study, we wanted to evaluate the differences in the response to a CT-scan radiation dose of 20 mGy in two lymphoblastoid cell lines with different radiosensitivity.

**Materials and Methods:** Several parameters were studied: gene expression, DNA damage, and its repair (by analyzing gamma-H2AX foci, chromosome breaks, and sister chromatid exchange), as well as cell viability, proliferation, and death.

**Results:** After 20 mGy of IR, the radiosensitive (RS) cell line showed an increase in DNA damage, and higher cell proliferation and apoptosis, whereas the radioresistant (RR) cell line was insensitive to this LD. Interestingly, gene expression analysis showed a higher expression of an antioxidant gene in the RR cell line, which could be used by the cells as a protective mechanism. After a dose of 500 mGy, both cell lines were affected by IR but with significant differences. The RS cells presented an increase in DNA damage and apoptosis, but a decrease in cell proliferation and cell viability, as well as less antioxidant response.

**Conclusions:** A differential biological effect was observed between two cell lines with different radiosensitivity, and these differences are especially interesting after a CT scan dose. If this is confirmed by further studies, one could think that individuals with radiosensitivity-related genetic variants may be more vulnerable to long-term effects of IR, potentially increasing cancer risk after LD exposure.

## INTRODUCTION

In recent years, the use of computed tomography (CT) imaging in medicine has increased rapidly due to its value as a diagnostic technique (Dahal and Budoff 2019). In fact, a comparison between 2008 and 2020 showed a two-fold increase in the number of CT examinations (UNSCEAR 2021). Notably, the dose of ionizing radiation (IR) employed in such technique, even though being low dose (LD) (<100 mGy), is higher than in other diagnostic tools. This is of great concern regarding public health because it is known that exposure to IR has long-term effects on human health, and these effects can be present at LD (Pearce 2011; Hauptmann et al. 2020). The harmful effects of IR are mainly due to DNA damage, such as single-strand breaks, double-strand breaks (DSB), DNA base alterations, and DNA-DNA or DNA-protein cross-links, producing genomic instability (Shimura and Kojima 2018).

Several *in vitro* studies have evaluated the effect that exposure to LD within the range administered in medical imaging may have on genetic material from human cells. Some of these studies have detected molecular changes by examining the number of DSB, detected as radio-induced foci of the phosphorylated histone H2AX (γH2AX) (Rothkamm and Löbrich 2003; Löbrich et al. 2005; Zelensky et al. 2020; Kaatsch et al. 2021); chromosomal aberrations, caused by unrepaired or misrepaired DSB (M’kacher et al. 2003; Golfier et al. 2009; Roch-Lefèvre et al. 2016; Tewari et al. 2016); frequencies of micronuclei, indicating chromosome breakage or loss (Joshi et al. 2014; Tewari et al. 2016); or gene expression of stress-responsive genes (Amundson et al. 1999; Ding et al. 2005; Franco et al. 2005; Gruel et al. 2008; Knops et al. 2012; Nosel et al. 2013; Kaatsch et al. 2021).

*In vivo* studies have also demonstrated changes at DNA level in peripheral blood lymphocytes from individuals undergoing a CT examination. It has been observed an increase and a subsequent disappearance of γ-H2AX (Löbrich et al. 2005; Rothkamm et al. 2007; Grudzenski et al. 2009; Pathe et al. 2011; Beels et al. 2012; Halm et al. 2014; Vandevoorde et al. 2015) and the presence of chromosomal aberrations (M’kacher et al. 2003; Stephan et al. 2007; Abe et al. 2015; Kanagaraj et al. 2015; Khattab et al. 2017).

The DNA damage caused by such LD of irradiation can increase cancer risk, a late effect of radiation that has become an essential component of radiation protection (Ali et al. 2020) and that may occur even at ultra-low doses of radiation (below 5 mGy) (Shimura and Kojima 2018). In fact, studies carried out in the United Kingdom (Brenner et al. 2001; Pearce et al. 2012; De Gonzalez et al. 2016), Australia (Mathews et al. 2013), Taiwan (Huang et al. 2014), Netherlands (Meulepas et al. 2019), and South Korea (Lee et al. 2021), showed an increased incidence of different types of cancer after the exposure to CT-scans, being this risk higher in younger patients. Recently, a large-scale multinational study observed an increase in hematological cancer risk after CT radiation exposure in children, adolescents and young adults (Bosch de Basea Gomez et al. 2023).

Notably, not all individuals respond the same to an identical radiation dose (El-Nachef et al. 2021); interindividual differences have been observed after exposure to medium or high radiation doses, even among individuals not affected by rare genetic syndromes. One possible explanation would be that those who present slight alterations in cell cycle or deficiencies in apoptosis or DNA repair pathways, probably due to genetic variants, are more prone to suffer from radio-induced cancer or high sensitivity after radiotherapy (Hornhardt et al. 2014). To predict radiation side-effects, several biomarkers have been evaluated in cells/organisms with different responses to IR, such as γH2AX or 53BP1 foci (Olive and Banáth 2004; Löbrich et al. 2005; Hornhardt et al. 2014; Borràs-Fresneda et al. 2016; Todorovic et al. 2019), gene expression profiles (Bishay et al. 2001; Yang et al. 2013; Young et al. 2014; Borràs-Fresneda et al. 2016; Todorovic et al. 2019), frequency of micronuclei and nucleoplasmic bridges (Bishay et al. 2001), chromosome aberrations (Pantelias and Terzoudi 2011; Borràs-Fresneda et al. 2016), cell viability and cell death (Borràs-Fresneda et al. 2016; Todorovic et al. 2019), cell proliferation (Todorovic et al. 2019), changes in cell cycle (Todorovic et al. 2019), and DNA methylation level (Newman et al. 2014). Most of these studies have been performed to find the differential toxic effect that radiotherapy may produce in normal tissue. Nonetheless, there is a lack of studies evaluating differences in the response to LD of IR, even though the human population is typically exposed to such doses and not to high doses. Using an animal model, Snijders et al. (2012) demonstrated a differential transcriptional response to LD of IR between two mice strains with different susceptibility to radiation-induced. Recently, using human fibroblast cell lines from patients with different radiosensitivity/susceptibility, Devic et al. (2022) observed differences in DSB recognition and repair after a head or chest CT scan dose.

The present study aimed to determine the differences in the response to a CT-scan radiation dose of 20 mGy between two lymphoblastoid cell lines, one radiosensitive (RS) and one radioresistant (RR), analyzing their gene expression, DNA damage, DNA repair capacity, cell viability, cell proliferation, and cell death after irradiation. Results were compared to the effect after a 500 mGy dose and after sham-irradiation.

## MATERIALS AND METHODS

### Cell lines and culture

Two Epstein-Barr virus-immortalized (lymphoblastoid cell lines (LCLs) from lung cancer patients, were used in the present study: 4060-200 and 20037-200. Both cell lines were kindly donated by Dr. Maria Gomolka and Dr. Sabine Hornhardt from the German Federal Office for Radiation Protection (BfS). One cell line is considered RS (4060-200) and the other non-RS (20037-200), which will be named RR from here on. Their radiosensitivity was previously determined by WST-1 and Trypan-blue survival assays (Guertler et al. 2011). Moreover, our research group had previously observed that these cell lines have differences in their levels of DNA damage, DNA repair capacity, cell death, and transcriptional response after 1 and 2 Gy irradiation (Borràs-Fresneda et al. 2016).

The LCLs were grown in suspension at 37°C in a 5% CO2 atmosphere in RPMI-1640 medium supplemented with 15% fetal bovine serum, L-glutamine 2 mM and penicillin/streptomycin (100 U/mL and 100 mg/mL, respectively).

### Irradiation

The two LCL cultures were irradiated in exponential phase with a dose of 20 mGy to emulate a CT dose index, the dose for abdominal/pelvic CT examinations in adults (Hanu et al. 2019). Irradiation with a radiation dose of 500 mGy was used as a positive control and sham irradiation as a negative control. Cells were irradiated with 6 MV photon beams from a TrueBeam linear accelerator (Varian Medical Systems, California, USA) located at Hospital de la Santa Creu i Sant Pau, Barcelona. To ensure homogeneous irradiation, an isocentric setup with two opposed fields (0° and 180°) was employed. Samples were placed inside two holes drilled in a 20 cm X 20 cm polymethyl methacrylate (PMMA) phantom with 20 cm thickness in the direction of the beams to provide full electron equilibrium to the samples. Monitor Units were calculated to deliver the prescribed dose to the samples. The effect of the table couch was taken into account for dose calculation. LINAC radiation beams were daily checked by means of two independent systems: Daily QA3 (Sun Nuclear, Wisconsin, USA) and Machine Performance Check (Varian Medical Systems). Before irradiation, all samples were warmed up at 37°C and placed inside the holes of the PMMA phantom. All irradiations were at the same dose rate of 0.167 Gy·s^-1^.

### RNA extraction and sequencing

RNA extraction of three replicas of the irradiated and sham-irradiated RR and RS cell lines was carried out 24 h post-irradiation with the RNeasy Mini Kit (Qiagen, Hilden, Germany) according to the manufacturer’s instructions. In the period between irradiation and extraction, the RS and RR cell lines were kept at 37°C in a 5% CO2 atmosphere. RNA concentration and purity were measured with a Nanodrop ND-1000 Spectrophotometer (Thermo Fisher Scientific, Waltham, MA, USA), and the integrity of the samples was assessed with the Agilent 2100 Bioanalyzer (Agilent Technologies, Santa Clara, CA, USA). All samples had an RNA Integrity number (RIN) higher than 6.60. RNA sequencing was carried out by the Centre for Genomic Regulation, Barcelona, Spain with the Illumina NextSeq 2000 (Illumina, San Diego, CA, USA). The read length sequenced was 2×125 bp, being 300 the maximum number of bases sequenced (300 cycles).

### Sequence Alignment, quantification, and differential expression analysis

The pre-processing of the reads was carried out with Trimmomatic (Bolger et al. 2014). The subsequent indexing of the human transcriptome (GRCh38) and quantification of the reads were performed with the software Salmon (Patro et al. 2017). To import the transcriptomic data and the metadata attachment into R, the R Bioconductor package Tximeta was used (Love et al. 2020).

The normalization of the data was achieved with the TMM method from the R Bioconductor package edgeR (Robinson et al. 2009). Data was then transformed to log2-counts per million (logCPM) and the mean-variance relationship was estimated using the function voom from the R Bioconductor package limma (Ritchie et al. 2015). The differential expression analysis was carried out with limma (Ritchie et al. 2015) removing unwanted variations with the Bioconductor R package sva (Leek JT, Johnson WE, Parker HS, Fertig EJ, Jaffe AE, Zhang Y, Storey JD 2022). P values were adjusted for multiple testing using the Benjamini-Hochberg approach to control the false discovery rate and p<0.05 was considered significant. Moreover, only genes with /log2FC/>0.85 were considered differentially expressed.

### Analysis of ***γ***-H2AX foci

In order to detect the radiation induction of DNA DSBs and their repair, analysis of γ-H2AX foci was performed. The kinetics of γ-H2AX foci following IR were obtained by performing an immunostaining of foci and microscope analysis at different time points after irradiation (0, 2, 4, and 20 h) following Borràs et al. (2016) procedures.

Automated slide scanning was done with Zeiss Axio Imager.Z2 microscope (Metasystems, Altlussheim, Germany) and a unique classifier were used to count foci in at least 100 cells for two replicas of each cell line and each experimental condition. A previous study with this classifier showed that the number of foci scored in 100 cells is enough to obtain a satisfactory result (Borràs et al. 2015). The γ-H2AX foci scoring was obtained with the MetaCyte software of the Metafer4 Slide Scanning System v3.10.2 (Metasystems). All signals were captured with a z-step size of 0.35 µm between 10 focal planes. The foci signals were captured using the SpOr filter.

### Analysis of chromosome gaps and breaks

After cell irradiation, three replicas of each cell line and condition were incubated for 24 h with 0.15 µg/mL of Colcemid (Gibco Thermofisher Scientific, Barcelona, Spain). The number of chromosome and chromatid gaps and breaks were obtained following Cabezas et. al. (2019) breakage assay procedures. Gaps were defined as a discontinuity shorter than the chromatid width or non-displacement.

### Sister chromatid exchange (SCE) assay

SCE are reciprocal exchanges of segments between chromatids and serve as a marker of chromosome instability. Their frequency after genotoxic conditions reflects DNA damage and the cell capacity to repair it by homologous recombination (Conrad et al. 2011). Three different replicas of the irradiated cell lines and the negative control were incubated at 37°C in a 5% CO_2_ atmosphere for 48 h with the thymine analog 5-bromo-2’-deoxyuridine (BrdU, Sigma-Aldrich, St. Louis, MI, USA). Colcemid was added 24 h before the extraction. The following steps until scoring the SCE values were the same used by Cabezas et al. (2019).

### Measure of cell proliferation

Proliferation and mitotic indexes were obtained for each cell line and condition as previously described by Cabezas et al. (2019). The proliferation index of three different replicas was obtained by counting the number of metaphase cells from the first, second, and third cell cycle (MI, MII, and MIII). The mitotic index of six replicas was calculated as the ratio of the number of mitotic cells in 1000 stimulated nuclei. Results were then normalized to sham-irradiated cells obtaining a FC increase.

### Cell death assay

Cell cultures of the irradiated and sham-irradiated RR and RS cell lines were kept at 37°C in a 5% CO2 atmosphere for 24 h after irradiation. Cells were then stained with the Annexin-V-FLUOS Staining Kit (Roche, Basel, Switzerland) following the manufacturer’s procedures. Results were measured by flow cytometry with CytoFLEX (Beckman Coulter Life Science, Pasadena, CA, USA), and analyzed with the software FlowJo v10 (BD Biosciences, Franklin Lakes, NJ, USA). To obtain the percentage of cell death, Annexin V positive cells (early and late apoptotic cells) were counted out of more than 20000 cells in six replicates of each experimental condition. Results were then normalized to sham-irradiated cells obtaining the FC induction of apoptosis.

### MTT assay for cell viability assessment

Cell viability for two replicas and both cell lines was determined at different time points after irradiation (24, 48, and 72 h) using the MTT assay. For the analysis, cells were transferred to a 96-well plate at a concentration of 8×104 cells/mL. Then, 10 µL of MTT (Sigma-Aldrich) were added to each well at the mentioned time points at a 5 mg/mL concentration for 1 h. After this, cells were incubated for 1 h with 100 µL of HCl 0.01 M to stop the reaction. The absorbances were then measured at a wavelength of 595 nm with the Sunrise absorbance microplate reader (TECAN, Männedorf, canton of Zürich, Switzerland). Background values, using only medium, were subtracted and results were normalized to sham-irradiated cells.

### Statistical analysis

The Kolmogorov-Smirnov test with Lilliefors correction was used to check for normality. In the case of normality compliance, the ANOVA test was chosen for the analysis and then multiple pairwise comparisons were performed with the Tukey test. For the rest of the analyses, the non-parametric Mann-Whitney test was used. All the statistical analyses were performed with R 4.2.1 software.

## RESULTS

### Differential expression analysis

The differential gene expression analysis of the cell lines was performed by extracting the RNA 24 h post-irradiation and comparing the gene expression profile of each cell line after irradiation with the one of sham-irradiated cells. The transcriptional response after irradiation at 20 mGy differed between the RS and the RR cell lines, and so did after 500 mGy irradiation. Table 1 presents the list of upregulated and downregulated genes for each cell line and dose grouped by their function; and Figure 1 shows the Volcano plot of the differential expression analysis. Upregulated genes were involved in DNA repair, cell cycle arrest, stress response, and antioxidant response, whereas downregulated genes were involved in inflammation, apoptosis, and cell survival. Some genes were upregulated (*HMOX1*) or downregulated (*GBP5*) in both cell lines and both doses, even though the FC increase in *HMOX1* expression was much higher in RR cells compared to RS cells (1.9 vs. 0.9 in both doses). Moreover, some genes seemed to be cell line specific, i.e. *RGS2* and *NRIP* were only downregulated in the RS cell line both after 20 and 500 mGy.

**Figure 1.**
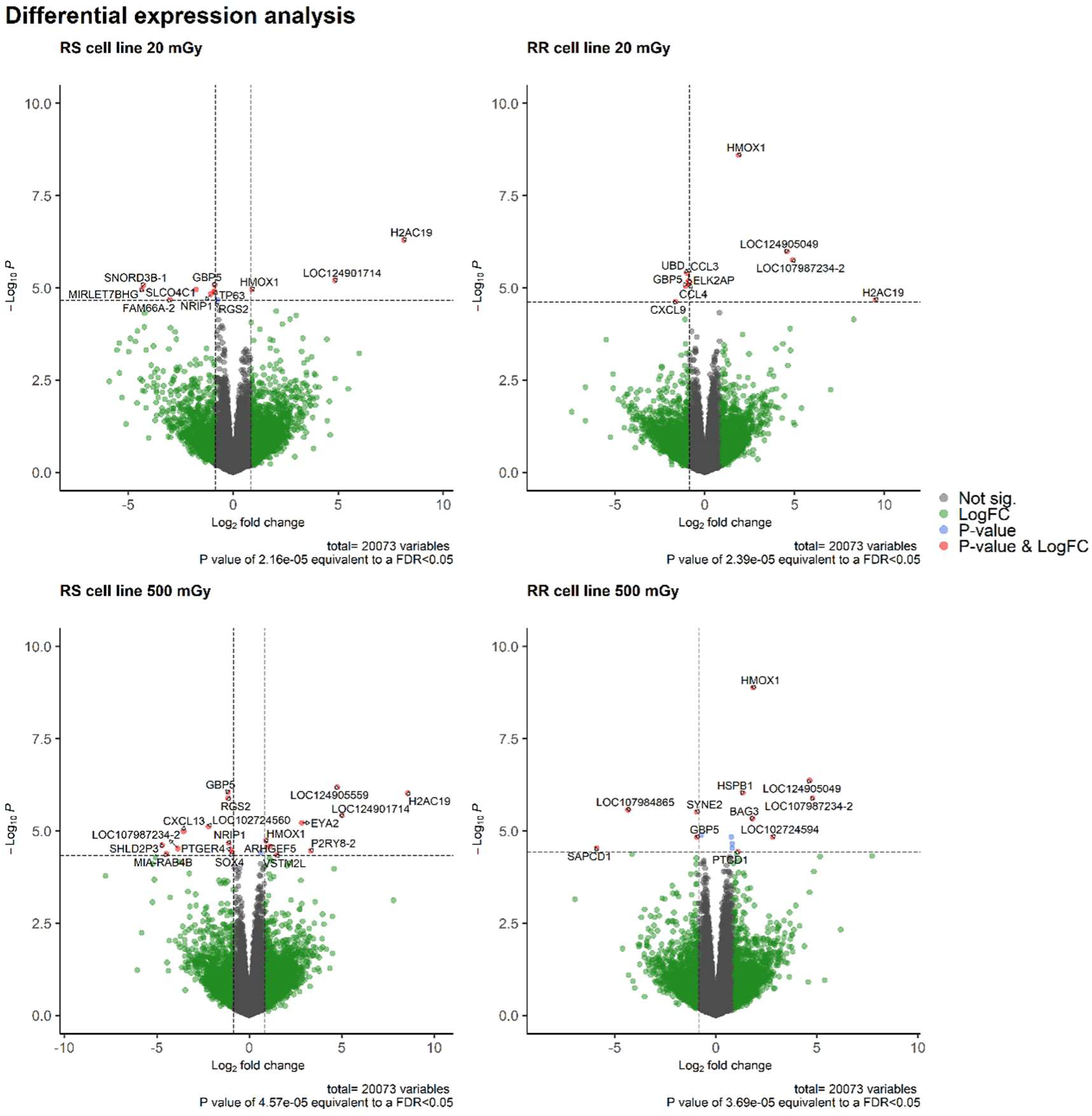
Differential gene expression analysis in the RS and the RR cell lines after irradiation compared to sham-irradiated cells. The horizontal dashed line marks the adjusted P value cut off (0.05) and the vertical one the /logFC/ cut off (0.85). The genes analyzed are represented as dots, and significant genes have their names by the dot. Not sig, Non-significant genes.

**Table 1.** Differentially expressed genes in the RR and RS cell lines after irradiation at 20 mGy and 500 mGy compared to sham-irradiated cells. Genes were classified according to their expression (upregulated and downregulated) and function.

|  | 20 mGy |  | 500 mGy |  | References |
| --- | --- | --- | --- | --- | --- |
|  | RS | RR | RS | RR |  |
| **Upregulated** |  |  |  |  |  |
| DNA repair |  |  | *EYA2* | *HSPB1* | (Krishnan et al. 2009; Zhou et al. 2018) (Arrigo 2017) |
| Antioxidant response | *HMOX1* | *HMOX1* | *HMOX1* | *HMOX1* | (Bao et al. 2016) |
| Stress Response |  |  |  | *BAG3* | (Sherman and Gabai 2022) |
| Others | *H2AC19*<br>*LOC124901714* | *H2AC19*<br>*LOC107987234-2*<br>*LOC124905049* | *H2AC19*<br>*LOC124901714*<br>*LOC124905559*<br>*ARHGEF5*<br>*P2RY8-2*<br>*VSTM2L* | *PTCD1*<br>*LOC107987234-2*<br>*LOC124905049*<br>*LOC102724594* |  |
| **Downregulated** |  |  |  |  |  |
| Inflammation | *GBP5* | *GBP5*<br>*UBD*<br>*CCL3*<br>*CCL4*<br>*CXCL9* | *GBP5*<br>*CXCL13* | *GBP5* | (Shenoy et al. 2012) (Kawamoto et al. 2019) (Bhavsar et al. 2015) (Griffith et al. 2014) (Griffith et al. 2014) (Griffith et al. 2014; Kim et al. 2019) |
| Apoptosis | *TP63* |  |  |  | (Suh et al. 2006) |
| Cell Survival | *NRIP1* |  | *NRIP1* |  | (Luan et al. 2016) |
| Others | *FAM66A-2*<br>*SLCO4C1*<br>*SNORD3B-1*<br>*MIRLET7BHG*<br>*RGS2* |  | *SHLD2P3*<br>*LOC102724560*<br>*LOC107987234-2*<br>*MIA-RAB4B*<br>*RGS2*<br>*PTGER4*<br>*SOX4* | *LOC107984865*<br>*SAPCD1*<br>*SYNE2* |  |

### Analysis of ***γ***-H2AX foci

The number of γ-H2AX foci was measured before irradiation and at 2, 4, and 20 h after 20 mGy and 500 mGy irradiation to see whether there were differences between the RS and the RR cell lines (Fig.2). After 20 mGy, the RS cell line showed a small increase in the number of foci, reaching its maximum at 2 h (2.26 foci/cell). Significant differences compared to background levels were only observed at 2 h post-irradiation (p<0.001, ANOVA), whereas a tendency could also be observed at 4 h (p=0.079, ANOVA). At 20 h post-irradiation the RS cell line had completely repaired the damage showing similar foci number than at time 0. On the other hand, no differences were observed between the number of foci at the different post-irradiation time points and the basal level in the RR cell line. Significant differences between cell lines were observed at 2 h after irradiation (p<0.001, ANOVA).

**Figure 2.**
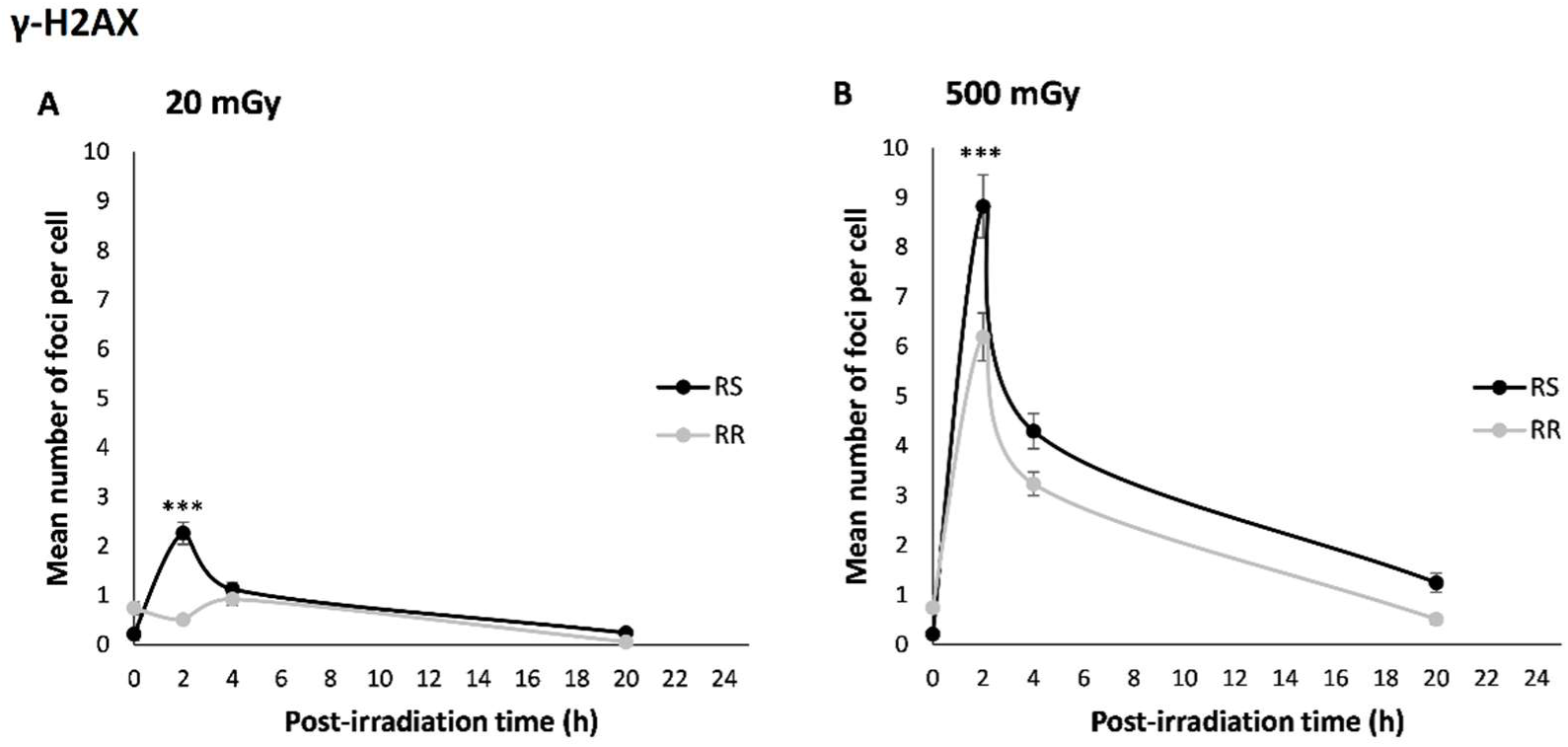
γ-H2AX kinetics in the RS and RR cell lines after irradiation at 20 mGy (A) and 500 mGy (B). Foci were scored at 2, 4, and 20 h after irradiation in more than 100 cells for each experimental condition. The basal level of γ-H2AX foci is represented as time 0. Data is plotted as mean ± SEM and results are representative of two independent experiments. Asterisks represent significant differences between the RS and the RR cell lines (***, p<0.001).

After a radiation dose of 500 mGy (Fig. 2B), the maximum number of γ-H2AX foci for both cell lines were observed 2 h post-irradiation but the number of foci counts differed, showing the RS cell line higher counts (8.82 foci/cell). The RS cell line showed differences at 2 h and 4 h compared to time 0 (p<0.001, ANOVA), and despite not being significant, also small differences could be observed after 20 h (p=0.11, ANOVA). As for the RR cell line, differences could also be observed at 2 h and 4 h post-irradiation (p<0.001, ANOVA), returning to its basal levels at 20 h post-irradiation. Significant differences between cell lines were only observed at 2 h post irradiation (p<0.001, ANOVA), and a tendency could also be observed at 4 h (p=0.072, ANOVA).

### Analysis of chromosome gaps and breaks

The number of chromatid and chromosome gaps and breaks was obtained 24 h post-irradiation (Fig. 3). After irradiation with 20 mGy, the RS cell line showed a tendency to present a higher number of gaps than the RR cell line (p=0.076, ANOVA), whereas after 500 mGy it showed a higher number of breaks (p=0.032, ANOVA). Overall, when counting both gaps and breaks the RS cell line showed a tendency to have more chromosome alterations per cell than the RR cell line after irradiation with 20 and 500 mGy. The RS cell line showed a slight increase in the number of alterations per cell compared to sham-irradiated cells in a dose-dependent manner, whereas RR cells showed the same number of alterations as sham-irradiated cells after 20 mGy and a slight increase after 500 mGy. However, these differences were non-significant.

**Figure 3.**
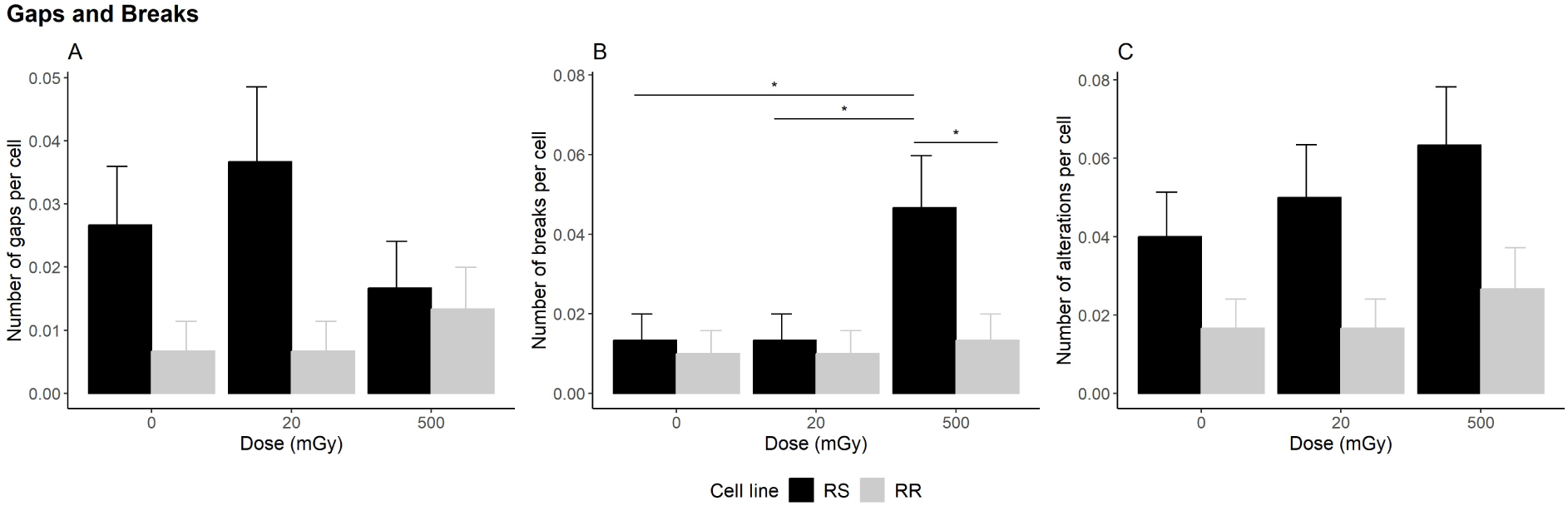
Number of chromosome and chromatid gaps (A) and breaks (B) and total aberrations per cell (C) in the RS and RR cell lines after irradiation at 20 mGy and 500 mGy and in sham-irradiated cells. Data from 300 cells are plotted as mean ± SEM. Asterisks represent significant differences (*, p<0.05).

### SCE assay

The number of SCE was measured 48 h post-irradiation in both cell lines (Fig. 4). The RS cell line presented a higher number of SCE than the RR cell line at 20 mGy and at 500 mGy (p<0.001, ANOVA). The RS cell line also showed differences compared to sham-irradiated cells, both after 20 mGy irradiation (p=0.01, ANOVA) and 500 mGy (p<0.001, ANOVA), whereas the RR showed a tendency to have less SCE per cell compared to sham-irradiated cells after 20 mGy (p=0.052, ANOVA), but it did show more SCE per cell than sham-irradiated cells after 500 mGy (p<0.001, ANOVA).

**Figure 4.**
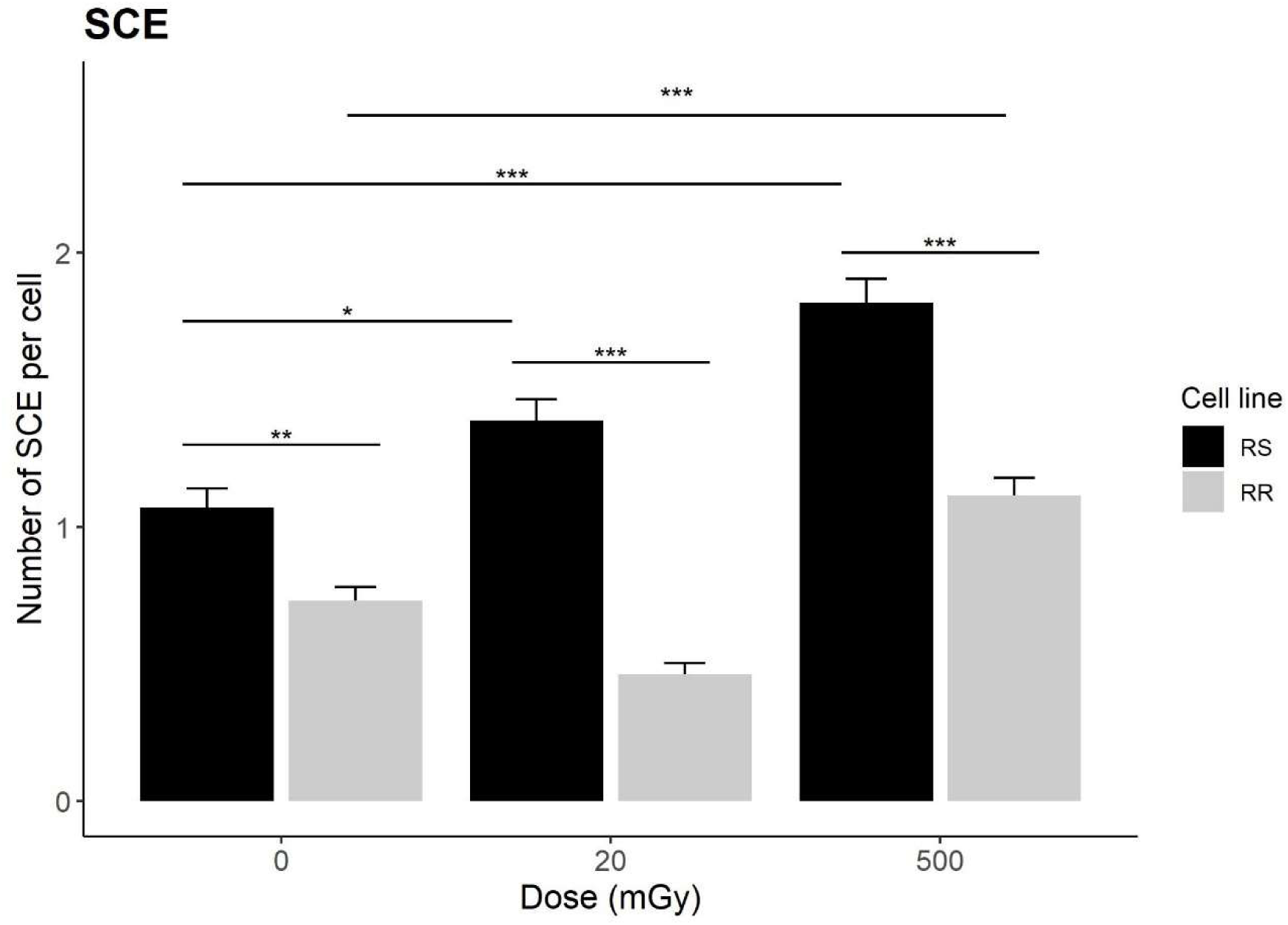
Mean number of SCE per cell in the RS and RR cell lines after sham-irradiation (0 mGy), and after irradiation at 20 mGy and at 500 mGy. Data from 300 cells are plotted as mean ± SEM. Asterisks represent significant differences (*, p<0.05; **, p<0.01; ***, p<0.001).

### Measure of cell proliferation

The mitotic index was evaluated 24 h after the irradiation at 20 mGy and 500 mGy and results are represented in Fig. 5. The RS cell line presented a higher mitotic index than the RR cell line in both cases; however, the difference between cell lines was not statistically significant (p=0.247 and p=0.082, respectively, Mann-Whitney). At 20 mGy the RS cell line showed a 1.5-fold increase compared to sham-irradiated cells (p= 0.049, Mann-Whitney), whereas the RR cell line did not change its mitotic index (p= 0.656, Mann-Whitney). After irradiation at 500 mGy, the RS cell line presented similar levels as sham-irradiated cells (p= 0.347, Mann-Whitney), while the RR showed a decrease in the mitotic index (p=0.007, Mann-Whitney).

**Figure 5.**
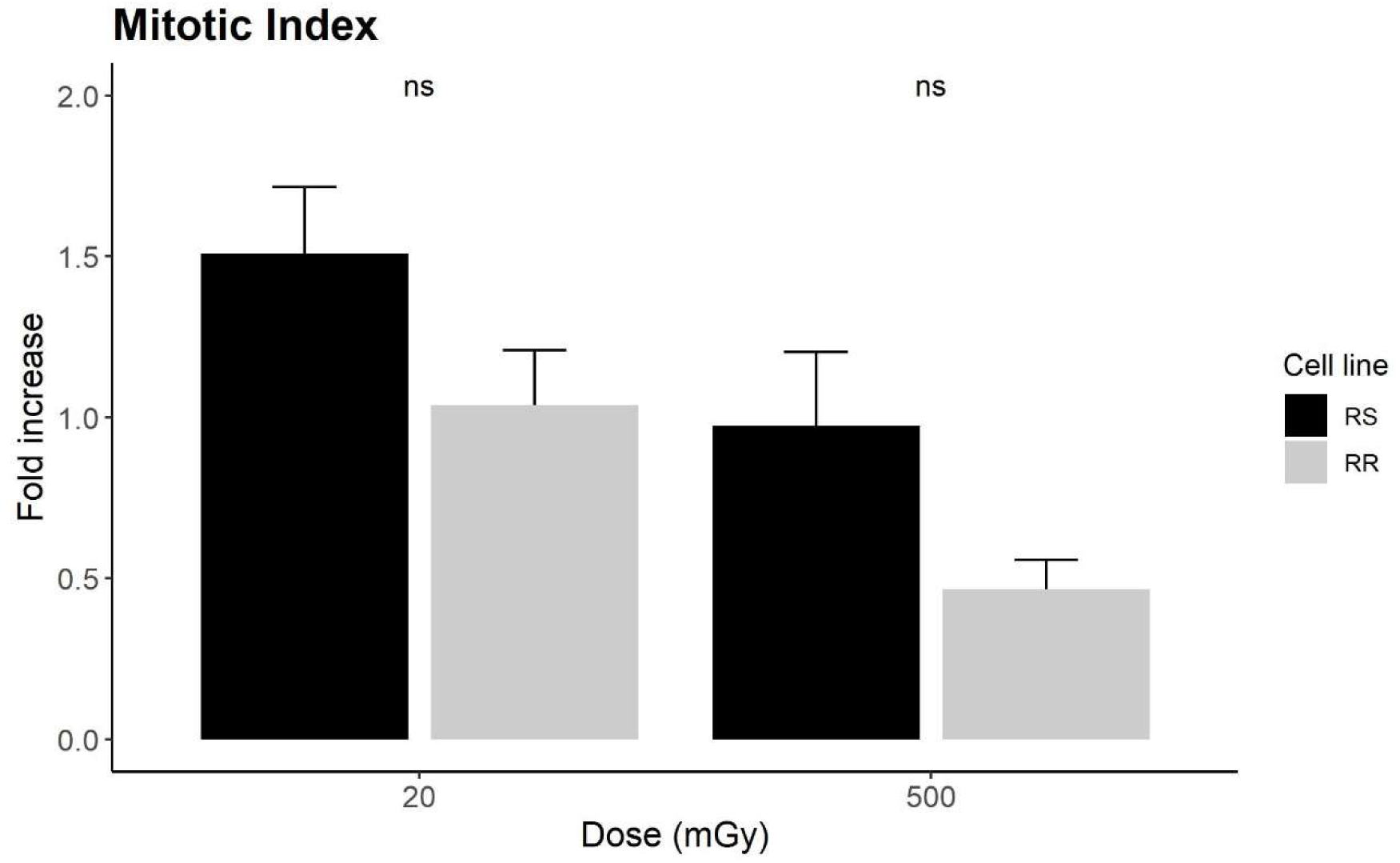
Excess of mitotic cells in the RS and RR cell lines after irradiation at 20 mGy or 500 mGy compared to the sham-irradiated cells. The mitotic index was measured 24 h post-irradiation. Data of six independent experiments are plotted as mean ± SEM.

As for the proliferation index, it was measured 48 h post-irradiation (Fig. 6). After 20 mGy irradiation, an increased number of cells at MI seemed to exist in the RR cell line compared to the RS one, whereas the RS cell line appeared to present more cells at MII and MIII. However, differences were not statistically significant (p=0.4 and p=0.2, respectively, Mann-Whitney). At 500 mGy, the RS cell line presented a reversed situation, with an increased number of cells at MI and a reduction of cells at MII and MIII compared to the RR cell line.

**Figure 6.**
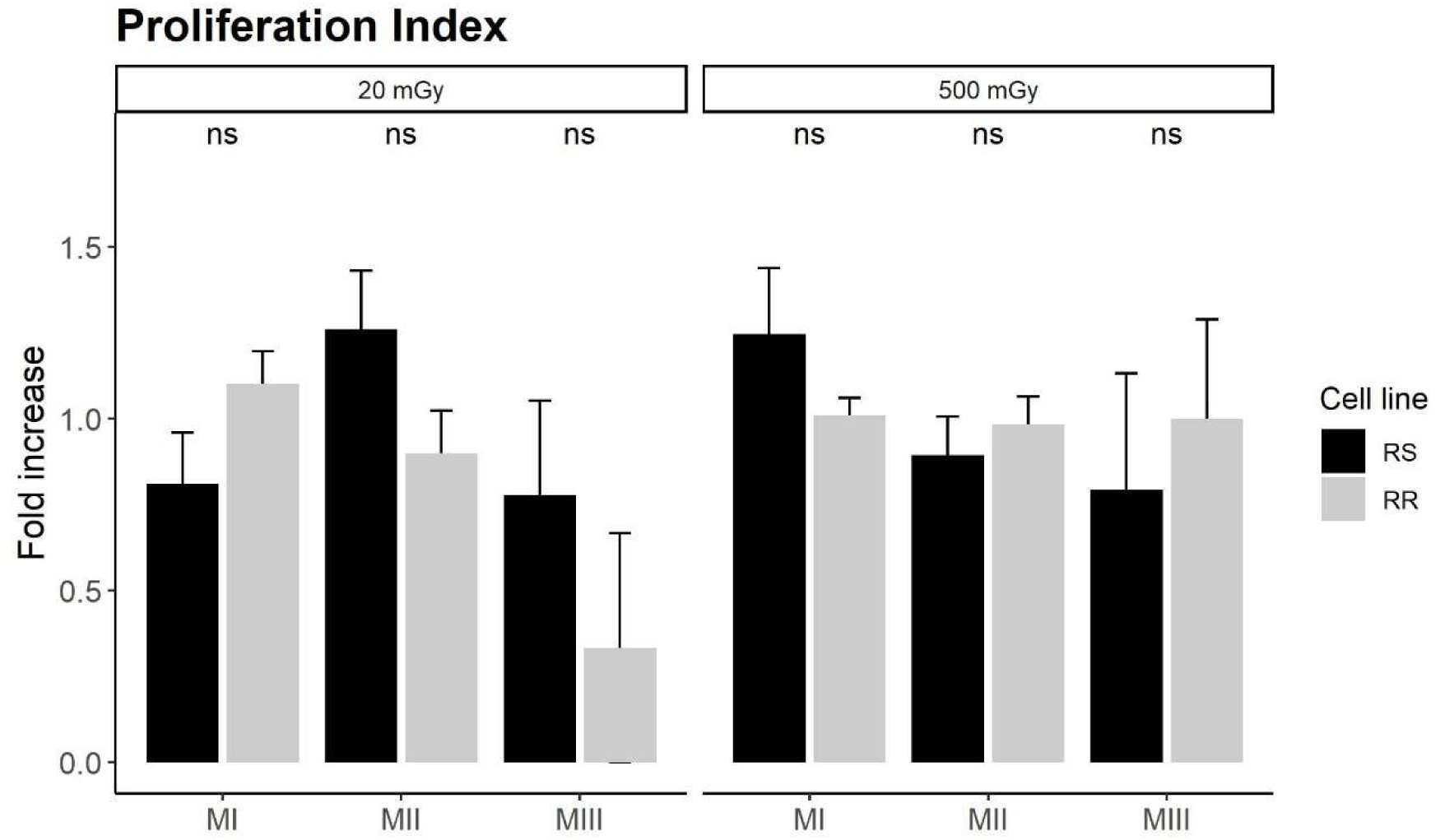
Fold increase of the number of cells at MI, MII, MII after irradiation with 20 mGy and 500 mGy compared to sham-irradiated cells for the RS and RR cell line. The proliferation state was measured 48 h post-irradiation. Data of three independent experiments are plotted as mean ± SEM.

### Cell death assay

The differences in the mortality induction between the RS and the RR cell lines were observed 24 h after irradiation at 20 mGy and 500 mGy (Fig. 7). The results showed an excess in the percentage of cell death in the RS cell line at 20 mGy (p=0.001, ANOVA) and at 500 mGy (p=0.002, ANOVA), compared to the RR cell line. Moreover, the RS cell line showed more apoptosis than sham-irradiated cells both after 20 mGy (p= 0.032, ANOVA) and after 500 mGy irradiation (p= 0.000, ANOVA), whereas the RR cell line showed very similar values of apoptosis than sham-irradiated cells after the dose of 20 mGy (p= 1, ANOVA) and a non-significant slight increase after the dose of 500 mGy (p= 0.365, ANOVA).

**Figure 7.**
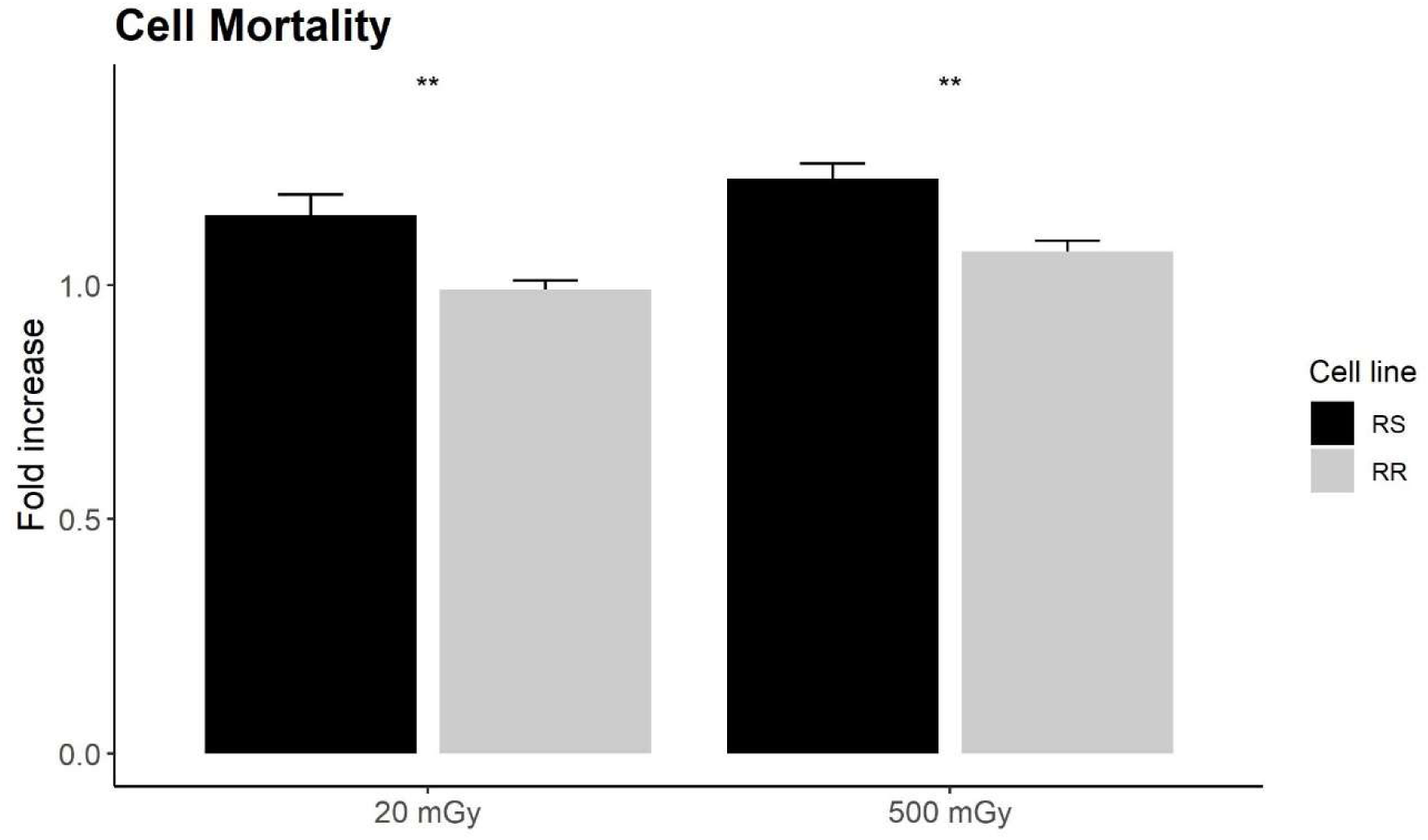
Excess of cell death in the RS and RR cell lines after irradiation at 20 mGy or 500 mGy compared to the sham-irradiated cell lines. Cell death was measured 24 h post-irradiation. Data from six independent experiments are plotted as mean ± SEM. Asterisks represent significant differences (*, p<0.05; **, p<0.01).

### Cell viability assessment

The results obtained for cell viability of the RS and RR cell lines at 24, 48, and 72 h after 20 mGy and 500 mGy irradiation can be seen in Fig.8. They were obtained from two different replicas and corrected by the cell viability of sham-irradiated cells. The percentage of viable cells did not differ between cell lines after 20 mGy irradiation at the three time points analyzed (p= 0.667, p= 1, p= 1, respectively, Mann-Whitney) and was similar to cell viability in sham-irradiated cells. However, differences between cell lines seemed to exist after 500 mGy irradiation, despite not being significant. The RR cells had a viability similar to sham-irradiated cells, whereas the percentage of viable cells was lower in the RS cell line for all time points (p= 0.333, p= 0.333, p= 0.333, respectively. Mann-Whitney).

**Figure 8.**
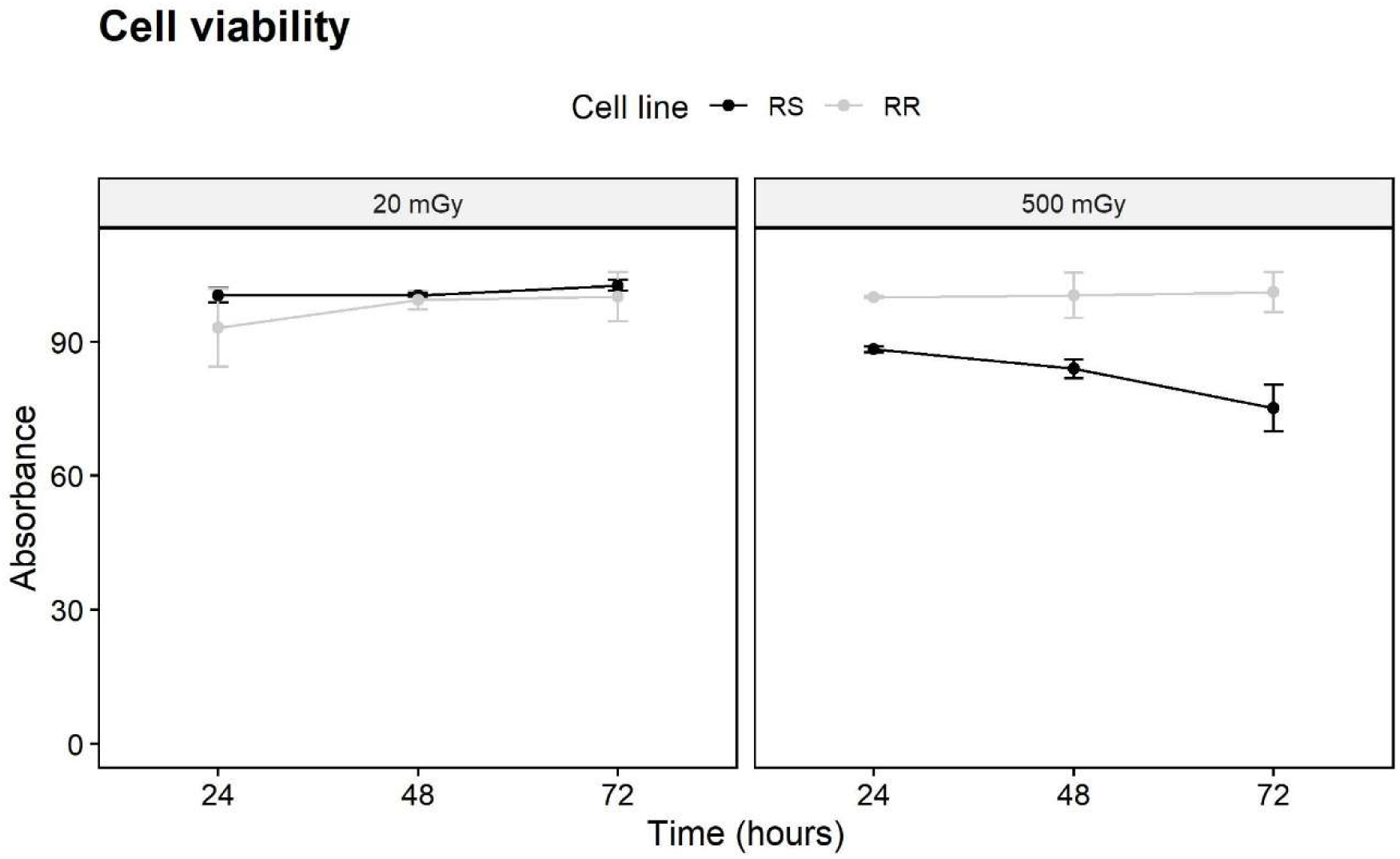
Percentage of cell viability in the RS and RR cell lines after irradiation with 20 mGy (left) or 500 mGy (right). Results were normalized to sham-irradiated cells. Data from two independent experiments are plotted as mean ± SEM.

## DISCUSSION

Several studies have detected an interindividual variation in radiosensitivity, mainly in individuals receiving radiotherapy (Andreassen et al. 2002; Barnett et al. 2009; Barnett et al. 2015). Most of these studies have been performed at medium or high IR doses, leaving behind LD, such as the ones employed in ever-growing medical imaging. In the present work, we wanted to evaluate if variation exists after exposure to a dose similar to a CT-scan, by using cell lines with different radiosensitivity as a model. Different parameters were analyzed: gene expression, DNA damage, and repair, as well as cell viability, proliferation, and death after irradiation at 20 mGy. Results were compared with those after a medium dose of 500 mGy. A summary of all the assays is presented in Table 2.

**Table 2.**
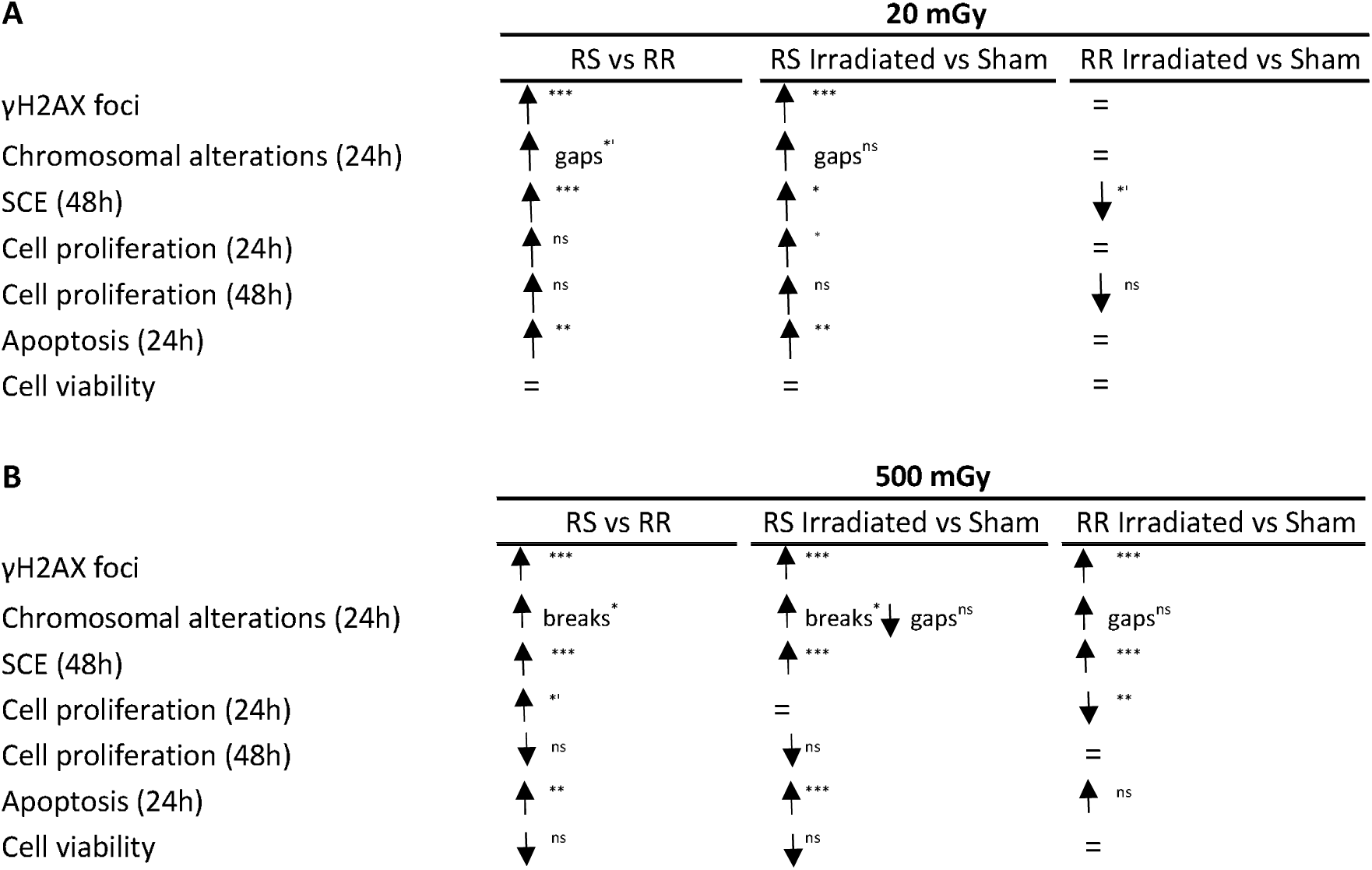
Summary of results obtained with all methods tested to compare RS and RR cells. Asterisks represent significant differences (*’, p<0.1; *, p<0.05; **, p<0.01; ***, p<0.001). Ns, non-significant differences.

After a LD of IR, RR cells did not present more DNA damage (γH2AX foci, chromosome alterations or SCE) nor changes in cell proliferation, cell death, or cell viability compared to sham-irradiated cells. Therefore, RR cells were insensitive to the CT scan dose. In contrast, RS cells showed a slight but significant increase in DNA DSB, measured as the number of foci per cell, compared to sham-irradiated cells. It has been reported that doses as low as 1 mGy can induce detectable γH2AX foci (Rothkamm and Löbrich 2003). However, according to our results, this would be only valid in RS cells but not in RR cells, in agreement with a recent study by Devic et al. (2022), who observed a great variability of foci induction and repair after head CT scan dose in cell lines with different levels of radiosensitivity. Concerning chromosome alterations, RS cells showed more gaps but the same number of breaks as RR cells and sham-irradiated cells. In these cells, an irradiation with 500 mGy was necessary to induce chromosome and chromatid breaks (Table 2B). It is believed that breaks may have a distinct biological effect than chromosome gaps (Oostra et al. 2012). Accordingly, a previous study has shown that the RS severely combined immunodeficiency (SCID/J) mice present this type of breaks after 50 mGy irradiation (Rithidech et al. 2013).

In both cell lines, the 20 mGy dose induced significant expression of the antioxidant gene *HMOX1*, in agreement with results obtained by other authors at 50 mGy (Bao et al. 2016). Interestingly, RR cells expressed double the amount of *HMOX1* compared to the RS cells. It is generally admitted that reactive oxygen species (ROS) can be elicited by LD IR (Tang et al. 2017) and that they can indirectly cause up to 80% of DNA damage in cells (Barry Halliwell and John M. C. Gutteridge 1999). The increase of the antioxidant HMOX1 levels would be used by the cell as a protective mechanism and could explain the lack of DNA damage in RR cells after a dose of 20 mGy.

Moreover, after 20 mGy, RS cells presented a tendency to have more cell proliferation at 24 h and at 48 h, and they also had a significant increase in apoptosis, compared to RR cells. These results agree with the changes of gene expression observed: one gene related to cell death (*TP63*) and one related to cell survival (*NRIP1*) were downregulated in the RS cell line, whereas no genes with these functions were differentially expressed in the RR cells. Several studies have reported that LD of IR can stimulate cell proliferation and cell cycle progression in different cell types (reviewed by Khan and Wang 2022; G. Yang et al. 2016). In our study, only RS cells increased cell proliferation after irradiation with 20 mGy, whereas the RR cells seemed to slow down proliferation at 48 h, albeit at non-significant levels. Besides this, it has been suggested that 1-2 DSB/cell are sufficient to induce apoptosis, even though at very low levels (Barazzuol et al. 2019), probably to eliminate cells with DNA damage that have not been repaired (Rothkamm and Löbrich 2003). This would be the case in the RS cell line irradiated at 20 mGy, where a slight but significant increase in apoptosis was observed. A simultaneous increase in cell proliferation and cell death is a likely explanation for the fact that overall cell viability measured with MTT assay was the same in RS cells compared to sham-irradiated cells and RR cells. In a previous study (Liang et al. 2011), cell viability after LD of IR was analyzed also by MTT assay in rat mesenchymal stem cells irradiated at doses ranging from 20 mGy to 100 mGy X-rays. After 20 mGy, no differences were observed compared to non-irradiated cells, only after 75 mGy was an effect observed.

After a dose of 500 mGy, both cell lines presented significantly more DNA damage (foci and SCE) than sham-irradiated cells. As for chromosome alterations, RR cells had a tendency to have more gaps but no breaks, whereas the RS cells had a significant increase in chromosome breaks, compared to sham-irradiated cells. As previously mentioned, breaks represent more severe damage in chromosomes than gaps. Overall, the RS cell line showed more γH2AX foci, chromosome breaks, and SCE than the RR cell line after a dose of 500 mGy. This is congruent with previous studies comparing cells with different radiosensitivity, after irradiation with a high dose (Olive and Banáth 2004; Lynam-Lennon et al. 2010; Pantelias and Terzoudi 2011; Schwartz et al. 2011; Goodarzi and Jeggo 2012; Borràs-Fresneda et al. 2016; Todorovic et al. 2019). The rate of DSB repair, observed as the kinetics of foci loss, was very similar between both cell lines, even though the RS cell line would still present residual foci 24 h after irradiation if the same repair rate was assumed. This would explain why at 24 h RS cells expressed the DNA repair gene *EYA2*, which mediates the dephosphorylation of H2AX at Tyr142 and promotes efficient DNA repair (Krishnan et al. 2009).

The lower levels of DNA damage in RR cells could be explained by the upregulation of enzymes with antioxidant functions. Besides expressing more *HMOX1* than RS cells, they also presented high expression of genes with stress response properties, such as the heat shock protein *HSPB1* and the cochaperone *BAG3*. High *HSPB1* expression can reduce the amount of ROS and nitric oxide levels (Arrigo 2017). In turn, BAG3 physically links HSPB1 with heat shock protein 70 (Hsp70), (Rauch et al. 2017) which is a key component of redox homeostasis (Zhang et al. 2022).

After 500 mGy, RS cells had the same cell proliferation rate as sham-irradiated cells at 24 h, whereas RR cells had significantly lower proliferation, resulting in a net increase in cell proliferation for RS cells when compared to RR cells. In contrast, at 48 h cell proliferation slowed down in RS cells but was recovered in RR cells. This would suggest a transient proliferation arrest in RR cells at 24 h. This proliferation arrest was not observed after 20 mGy. Interestingly, other reports have observed a defined threshold for cell cycle arrest at 200 mGy (Barazzuol et al. 2019). The same authors reported an initial arrest and a posterior recovery by 48 h, similar to what we observed for RR cells. This arrest would serve to repair the DNA damage caused by radiation, with less need for induction of apoptosis to eliminate injured cells. Therefore, RR cells had no changes in cell viability after 500 mGy irradiation. It seems that RR cells would need a higher dose to have their viability compromised. On the other hand, RS cells had considerably more DNA damage than RR cells and consequently, more cell death and a tendency to less proliferation after 24 h, which resulted in a progressive net decrease in cell viability, although at non-significant levels. Interestingly, RS cells showed downregulation of the pro-survival gene *NRIP1* 24 h after irradiation.

The study of gene expression is of great interest to identify possible processes altered after exposure to IR. Remarkably, there was a downregulation of genes involved in the inflammatory immune response, especially in the RR cell line irradiated at 20 mGy. It is well known that LD of IR can induce an anti-inflammatory response, even though this effect has been usually observed at doses between 0.1 and 1 Gy (reviewed in Lumniczky et al. 2021). Notably, there is a connection between anti-inflammatory response and antioxidant response after LD of IR. It has been reported that IR doses smaller than 1 Gy activate the nuclear factor erythroid 2 (NRF2). NRF2 is a transcription factor that can downregulate the expression of pro-inflammatory molecules, and increase the expression of antioxidant genes, such as *HMOX*-1, which was found to be upregulated in the present study (Javadinia et al. 2021).

In a previous study, we analyzed the differences between the same cell lines used in the present study after 1 Gy and 2 Gy irradiation (Borràs-Fresneda et al. 2016). Contrary to what has been observed in the present study with lower doses, in that case, we observed that RS cells had a slower rate of γH2AX foci disappearance, which, would correspond to a different DNA repair capacity. This could suggest that radioresistance to LD (20 mGy) and intermediate doses (500 mGy) would rely on antioxidant defenses, whereas radioresistance to higher doses (1 Gy and 2 Gy) would depend more on DNA repair capacity. This divergent response between LD and high doses has been previously observed by other authors (Sampadi et al. 2022). In conclusion, we were able to observe differences between a RS and a RR cell line after an irradiation with a dose similar to a CT-scan dose. RR cells seemed to be insensitive to this dose, whereas RS cells showed DNA DSB, cell proliferation, and apoptosis, as well as less antioxidant response. Erroneous repair of DNA damage and the presence of oxidative stress can induce chromosome alterations, sequence mutations, and overall genome instability, which can contribute to carcinogenesis. If this is confirmed with further studies, one could think that individuals with genetic variants conferring radiosensitivity could be more affected by LD of IR than other individuals, and this might be related to cancer proneness after CT scan exposure.

In the field of radiological protection, a linear no-threshold model (LNT) is currently used to estimate the risk of cancer (and other stochastic effects) after LD of IR (reviewed by UNSCEAR 2021; Laurier et al. 2023). Accordingly, some recent epidemiological studies showed an increase in cancer mortality in nuclear industry workers (Richardson et al. 2023) or an increase in hematological cancer after a CT-scan (Bosch de Basea Gomez et al. 2023). However, other studies suggest other options, such as that the LNT model overestimates the risk of cancer after LD of IR, that there are different slopes of dose-response for LD and high doses, or even that there is a threshold below which no deleterious effects would exist (reviewed by Laurier et al. 2023). The present study shows that, after a CT-scan dose, there are genotoxic and molecular effects and that these effects are different depending on the radiosensitivity of the cells.

We want to thank Jéssica Martínez for her help in carrying out the experiments.

This project has received funding from the Euratom research and training program 2014-2018 under grant agreement No 755523.

## Data availability statement

Research data are stored in the institutional repository from UAB and will be shared upon request to the corresponding author.

